# A SARS-CoV-2 peptide vaccine which elicits T-cell responses in mice but does not protect against infection or disease

**DOI:** 10.1101/2022.02.22.481499

**Authors:** Victoria K. Baxter, Elizabeth J. Anderson, Sharon A. Taft-Benz, Kelly Olsen, Maria Sambade, Kaylee M. Gentry, Wolfgang Beck, Jason Garness, Allison Woods, Misha Fini, Brandon Carpenter, Christof C. Smith, Mark T. Heise, Benjamin Vincent, Alex Rubinsteyn

## Abstract

We vaccinated BALB/c mice with peptides derived from the SARS-CoV-2 proteome selected *in silico* to elicit T-cell responses and/or B-cell responses against linear epitopes. These peptides were administered in combination with either of two adjuvants, poly(I:C) and the STING agonist BI-1387466. Antibody responses against predicted linear epitopes were not observed but both adjuvants consistently elicited T-cell responses to the same peptides, which were primarily from the set chosen for predicted T-cell immunogenicity. The magnitude of T-cell responses was significantly higher with BI-1387466 compared with poly(I:C). Neither adjuvant group, however, provided any protection against infection with the murine adapted virus SARS-CoV-2-MA10 or from disease following infection. In light of more recent evidence for protection from severe disease mediated by CD8+ T-cells, we suspect that the epitopes selected for vaccination were not presented by infected murine cells.

## Introduction

Strong and persistent T-cell responses to SARS-CoV-2 have been demonstrated across many studies of convalescent patients (7–10). These responses have been thought to play an important, albeit secondary, role in viral clearance and have also been measured as secondary endpoints in many trials of vaccines against SARS-CoV-2 (11–13). While CD4+ T-cell responses have a well understood role in the promotion of antibody responses(14), it is less clear whether either CD4+ or CD8+ T-cell responses independent of B-cells provide protection against infection with SARS-CoV-2 or attenuate disease severity. The small body of early studies on this issue reported contradictory results which varied across animal models (15–18). As the SARS-CoV-2 pandemic continued, however, it became clear that T-cell responses even in the absence of antibody neutralization play some kind of protective role in humans (2, 3). More definitively, several T-cell directed vaccines were able to demonstrate protection from infection and disease mediated by specific T-cell responses in the absence of neutralizing antibodies (1, 5, 6).

Of the early studies evaluating the role of SARS-CoV-2 specific T-cells in viral clearance the most surprising was *Hasenkrug et al*. (15), which showed that depleting the majority of CD4+ and nearly all CD8+ T-cells in rhesus macaques did not significantly alter the course of infection or symptoms of disease upon reinfection. On the other hand, a similar study by McMahan et al.(16) found that CD8+ T-cells did play an assistive role in viral clearance which is most evident when antibodies are absent and/or insufficiently neutralizing. *Israelow et al(17)* examined the relative roles of humoral and cellular immunity from both prior infection and vaccination in K18-hACE2 by a combination of depletion and transfer studies. Their findings confirm that neutralizing antibodies provide protection even in the absence of CD8+ T-cells. Conversely, transfer of SARS-CoV-2 specific CD8+ T-cells to Rag1 deficient mice, in the absence of humoral immunity, provided noticeable but incomplete protection from infection. A complementary study of nucleocapsid vaccination in Syrian hamsters and K18-hACE2 mice by *Matchett et al.(18)* showed T-cell mediated partial protection from severe disease.

The confusing and contradictory results of early SARS-CoV-2 T-cell studies motivated the work in this study. We would like to highlight, however, that more recent studies have clearly demonstrated the protective capacity of T-cells during SARS-CoV-2 infection. *Pardieck et al (1)* showed that synthetic long peptide vaccination using the spike epitope S539-546 (VNFNFNGL) protects K18-hACE2 mice from lethal infection. SImilarly, *Carter et al (4)* demonstrated that a T-cell targeting mRNA vaccine containing 11 conserved SARS-CoV-2 epitopes significantly reduces lung viral titers a week after infection. Lastly, *Arieta et al (6)* showed that a version of the BioNTech mRNA vaccine containing non-Spike antigens is protective against disease and infection in Syrian golden hamsters. Taken together with studies of robust T-cell responses in humans against diverse SARS-CoV-2 variants (2, 19), these studies firmly establish T-cells as a potent second line of defense against SARS-CoV-2 infection.

Vaccines using synthetic peptides as antigens have a long and often unsuccessful history of use against pathogens. Peptide vaccines have been designed to target linear B-cell epitopes for pathogens such as FMDV (20), DENV (21, 22), and HIV (23) but are not typically are able to elicit significant numbers of neutralizing antibodies except in rare cases where a target a highly functional conserved linear epitope can be found on the surface of a viral receptor. Furthermore, antigenic peptides must either be conjugated to a carrier or are otherwise modified to improve valency and conformational stability (24, 25). Eliciting T-cell responses with peptide vaccination is more straight-forward, which has resulted in much interest in their use for therapeutic cancer vaccination (26, 27). In particular, adjuvants such as polyinosinic:polycytidylic acid and its derivative poly-ICLC have become commonly used for cancer vaccination in combination with synthetic peptides (28–31). There have been a few attempts at achieving protection from viral infection with T-cell directed peptide vaccines, such as for HIV (32) and EBOV (33), but experience with T-cell vaccines is much more limited than the rich body of work relating to achieving protective antibody responses.

In this study we use viral challenge following peptide vaccination to examine the potential for T-cell responses in the absence of neutralizing antibody responses to protect mice from infection with SARS-CoV-2-MA10 or disease subsequent to infection(34). The vaccine contains peptides selected for predicted T-cell immunogenicity, compared with a second group of peptides selected for antibody responses against linear epitopes. Peptide antigens are combined with either of two different adjuvants known for eliciting potent T-cell responses, poly(I:C)(35) and the STING agonist BI-1387466(36).

## Methods

### Vaccine peptide selection

Vaccine contents were selected in a previous publication (37), whose methods will be briefly summarized here. The protein sequences of the ancestral Wuhan-1 SARS-CoV-2 strain were analyzed for candidate linear B cell epitopes along with predicted human CD4+ and CD8+ T-cell epitopes which coincide with murine MHC-I/MHC-II ligands. B-cell epitope regions were chosen from linear epitope mapping studies of convalescent patient serum, followed by computational filtering for predicted surface accessibility, sequence conservation, spatial localization near annotated functional domains of the spike protein, and avoidance of glycosylation sites. T-cell epitopes were derived in a purely computational manner, starting with MHC binding predictions across a variety of high frequency HLA alleles. These predicted MHC ligands were further filtered by predicted immunogenicity, sequence conservation, and source protein abundance. 27mer vaccine peptides were selected to optimize the number of adjacent predicted T-cell epitopes, predicted murine MHC ligands for the H2-b and H2-d haplotypes, along with *in silico* prediction of manufacturability. The selection process ultimately yielded 22 candidate vaccine peptides, which were manually curated to 16 sequences in order to eliminate redundancy between highly overlapping sequencing.

### Mouse vaccination for immunogenicity study

All mouse work was performed according to IACUC guidelines under UNC IACUC protocol ID 20-121.0. Vaccine studies were performed using BALB/c mice with free access to food and water. Mice were ordered from Jackson Laboratories and vaccinated at 8 weeks of age. Equal numbers of male and female mice were used per group, vaccinated subcutaneously with poly(I:C) (Sigma-Aldrich cat. #P1530) or STING agonist BI-1387446 alone as controls or in combination with 16 synthesized vaccine peptides, consisting of 480 μg total peptide per vaccination (divided equimass per peptide, 30μgs each).

Control groups were n = 3 mice per group, and the peptide with adjuvant combinations were n = 6 mice per group. 50 μg of poly(I:C) was added to peptide mix or PBS control per vaccination. For the STING agonist BI-1387446 vaccinations, 10μgs was diluted in 200uls of PBS, and injected in 50uls quantities to form a 1 cm square. Peptide mix or PBS control was injected within this 1 cm square. Mice were vaccinated on days 1 and 8, cheek bleeds obtained on days 8 and 15, and sacrificed with cardiac bleeds performed on day 22.

### Peptide ELISA

Serum obtained from cardiac bleeds on day 21 and cheek bleeds on experimental days 7 and 14 were tested for antibody response to the predicted B cell peptide epitopes used for vaccinations via peptide ELISAs. Plates were coated with 5μg/mL of target peptide using coating reagent from the Takara Peptide Coating Kit (Takara cat. #MK100). Measles peptide was utilized as a negative control, and Flag peptide was also plated as an experimental control. Plates were blocked with a blocking buffer according to the manufacturer’s protocol. Serum was plated in duplicate wells with serial dilutions, and anti-FLAG antibody was plated in the experimental control wells. Rabbit anti-mouse IgG HRP (Abcam ab97046) was utilized as a secondary antibody. TMB substrate (Thermo Fisher Scientific cat. #34028) was added, development was stopped with TMB Stop solution (BioLegend cat. #423001), and plates were read at 450 nm.

### Protein ELISA

Serum obtained from cardiac bleeds on day 21 was utilized for ELISA testing for antibody response to SARS-CoV-2 spike (S) protein. Nunc Maxisorp plates (Thermo Fisher Scientific) were coated with S protein (generously provided by Ting Lab at UNC), or BSA as a negative control and incubated overnight. Plates were blocked with 10% FBS in PBS, washed, and serum plated in duplicate wells with serial dilutions. 6x His Tagged monoclonal antibody (Thermo Fisher Scientific) was also plated as an experimental control. Goat anti-mouse IgG HRP (Thermo Fisher Scientific) was added to washed plates as a secondary antibody. TMB substrate (Thermo Fisher Scientific) was added, development was stopped with TMB Stop solution (BioLegend), and plates were read at 450 nm.

### T-cell response quantification with ELISpot

Mice were sacrificed on day 21 and spleens collected. Spleens were mechanically dissociated using a GentleMACS Octo Dissociator (Miltenyi Biotec) and passed through a 70-μm filter. RBC lysis buffer (Gibco cat. #A1049201) was used to remove red blood cells. Cells were washed then passed through 40-μm filters. Splenocytes were counted and viability measured. 250,000 splenocytes were plated per well to perform an IFN-gamma ELISPOT per protocol using a Mouse IFN-gamma ELISPOT Kit (BD Biosciences; cat. #551083). Each sample was plated in triplicate with peptide or without peptide as a negative control. 30ug/ml of each peptide was used to stimulate appropriate T cells during a 72hr ELISPOT incubation. Plates were developed using AEC substrate kit per protocol (BD Biosciences; cat. # 551951). Activity values, representing the amount of signal per well, were determined using an AID Classic ERL07 ELISpot Reader. The average activity values of the negative controls were subtracted from all activity values of individual samples stimulated with peptide.

### Mouse vaccination for viral challenge

Using the same IACUC protocol described above, female BALB/c mice were used, 5 per group. The groups consisted of the 16 synthesized vaccine peptides with poly(I:C) (Sigma-Aldrich cat. #P1530), 16 synthesized vaccine peptides with STING agonist BI-1387446, and PBS control group. Each mouse was vaccinated subcutaneously with 30μgs of each peptide, and 50μgs of poly(I:C) or 10μgs of STING agonist. Mice were vaccinated on days 1 and 8, then transferred on day 22 for viral challenge assay.

### Live virus challenge

Mice were lightly anesthetized with 50 mg/kg ketamine along with 15 mg/kg xylazine and were then intranasally inoculated with 10^4 pfu of SARS-CoV-2-MA10(34) diluted in 50uL PBS.

### Body weight and clinical scoring

Body weights and clinical scores were recorded daily. Clinical disease was assessed using a 6-point scale(38): 0 = normal; 1 = piloerection; 2 = piloerection and kyphosis; 3 = piloerection, kyphosis, and reduced movement; 4 = piloerection, kyphosis, minimal spontaneous movement, +/-labored breathing (humane endpoint); 5 = moribund, dead, or euthanized. On 5 days post infection (DPI), mice were euthanized by an overdose of isoflurane anesthesia, and lungs were collected.

### Gross lung discoloration scoring

Gross lung discoloration scores were assigned as follows(38): 0 = normal, pink lungs; 1 = severe discoloration affecting less than 33% of the lung surface area or mild to moderate discoloration affecting less than 67% of the lung surface area; 2 = severe discoloration affecting 34% to 67% of the lung surface area or mild to moderate discoloration affecting 68% to 99% of the lung surface area; 3 = severe discoloration affecting 68% to 99% of the lung surface area or mild to moderate discoloration affecting 100% of the lung surface area; and 4 = severe discoloration affecting 100% of the lung surface area.

## Results

T-cells from mice vaccinated with either poly(I:C) or STING agonist as an adjuvant showed similar patterns of response but with significantly higher levels of activity for STING agonist (Figure 1). These responses were concentrated on peptides which had been selected for predicted T-cell immunogenicity, although one of six predicted B-cell targeting peptides also showed T-cell responses. Though we detected mild antibody responses to some of the peptides chosen for predicted linear B-cell epitopes (Figure 2), serum from vaccinated mice did not show meaningfully high levels of antibody binding to SARS-CoV-2 spike protein (Figure 3). This indicates that either B-cell responses were not elicited or the linear epitopes targeted by vaccine peptides are not a good match for the conformational structure of spike. Though no neutralization study was performed, we can infer that neutralization without spike binding antibodies is extremely unlikely. Challenge of vaccinated mice with a murine adapted strain (SARS-CoV-2-MA10) demonstrated that elicited T-cell responses did not confer protection from infection or disease (Figures 4 and 5). Specifically, vaccinated mice did not differ significantly from control mice with regards to clinical scoring (Figure 4A), change in body weight (Figure 4B), lung discoloration (Figure 4C), or lung viral titers (Figure 5).

**Figure 1.**
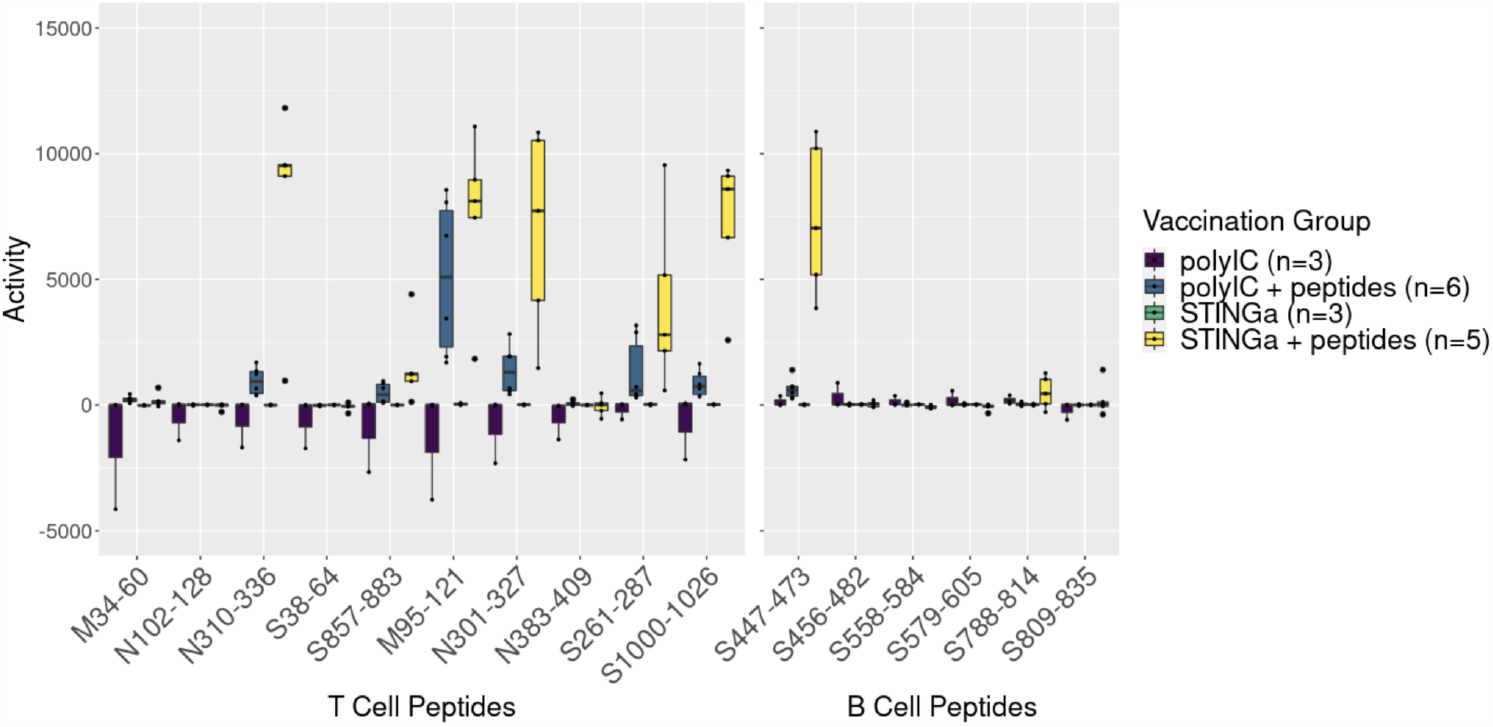
T-cell responses to individual vaccine peptides in BALB/c mice to vaccination with adjuvant only (polyIC, STINGa) and both adjuvants combined with peptides. Of the six peptides with noteworthy T-cell responses, five were included in the vaccine set for their predicted T-cell immunogenicity. The peptides with highest T-cell responses from STING agonist adjuvant also have the highest responses for the poly(I:C) group but poly(I:C) consistently attains order of magnitude lower responses.

**Figure 2.**
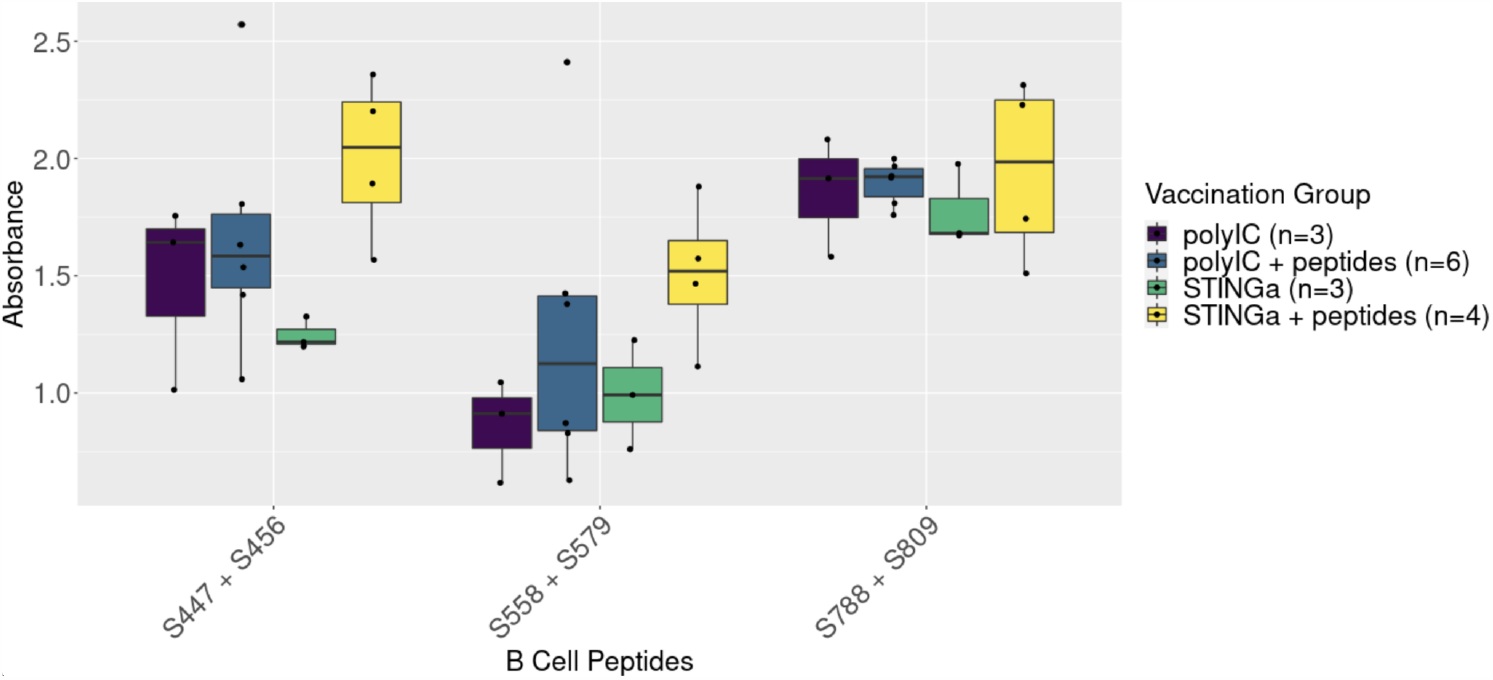
Antibody responses to pairs of vaccine peptides predicted to contain linear B-ecell epitopes in BALB/c mice after vaccination with adjuvant only (polyIC, STING) or adjuvant combined with peptides. While vaccination with poly(I:C) + peptides does not seem to meaningfully increase antibody responses against peptides, there is a meager increase over baseline when using the BI-1387446 STING agonist as an adjuvant, especially in the pair of peptides overlapping an RBD derived B-cell epitope (S447 + S456).

**Figure 3.**
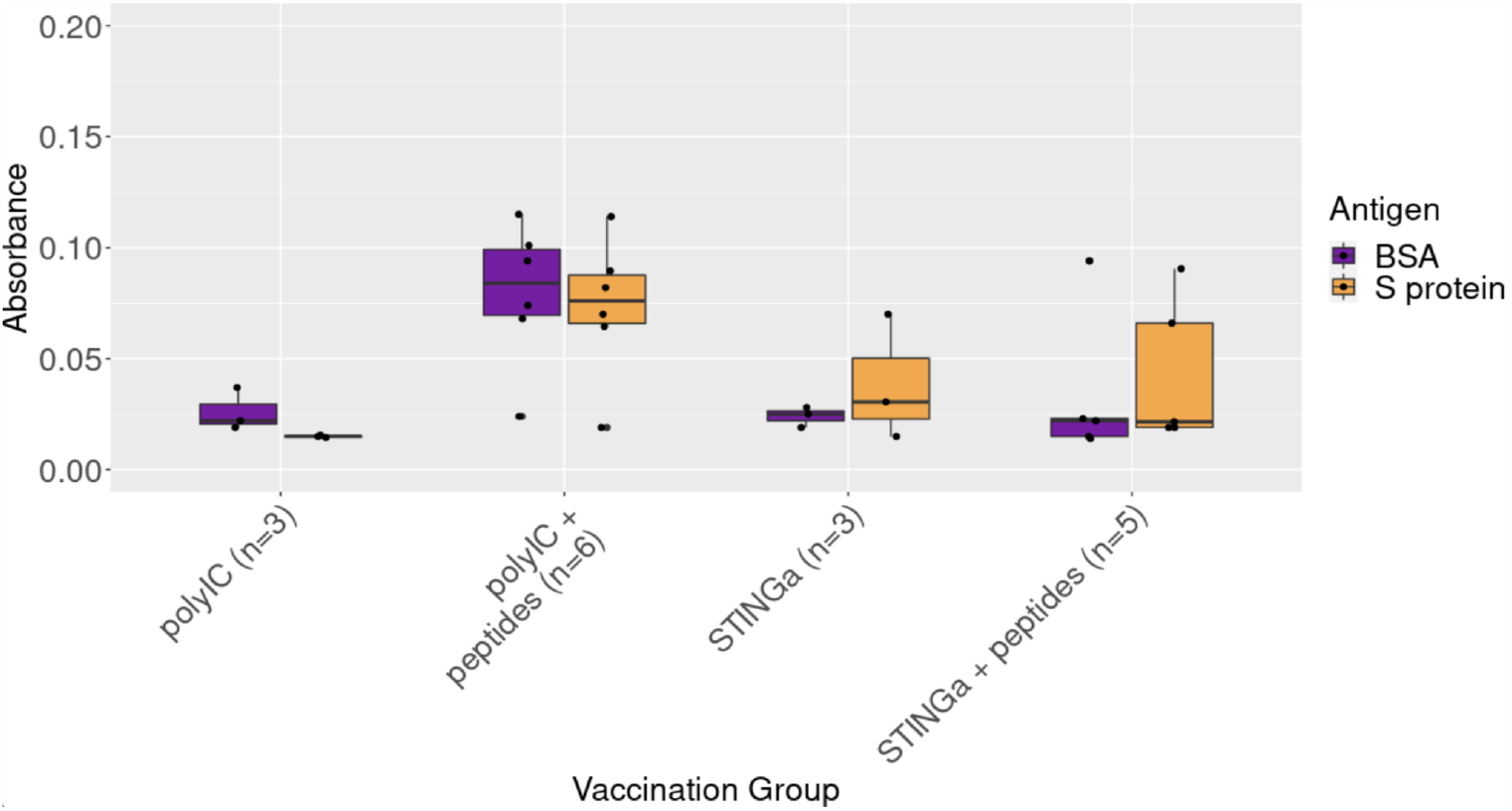
Antibody responses to SARS-CoV-2 spike protein in BALB/c mice after vaccination with adjuvant only (polyIC, STING) or adjuvant combined with peptides as measured by protein ELISA with either spike protein or a negative control (BSA). No group achieved significantly more antibody absorbance with spike protein as compared with BSA.

**Figure 4.**
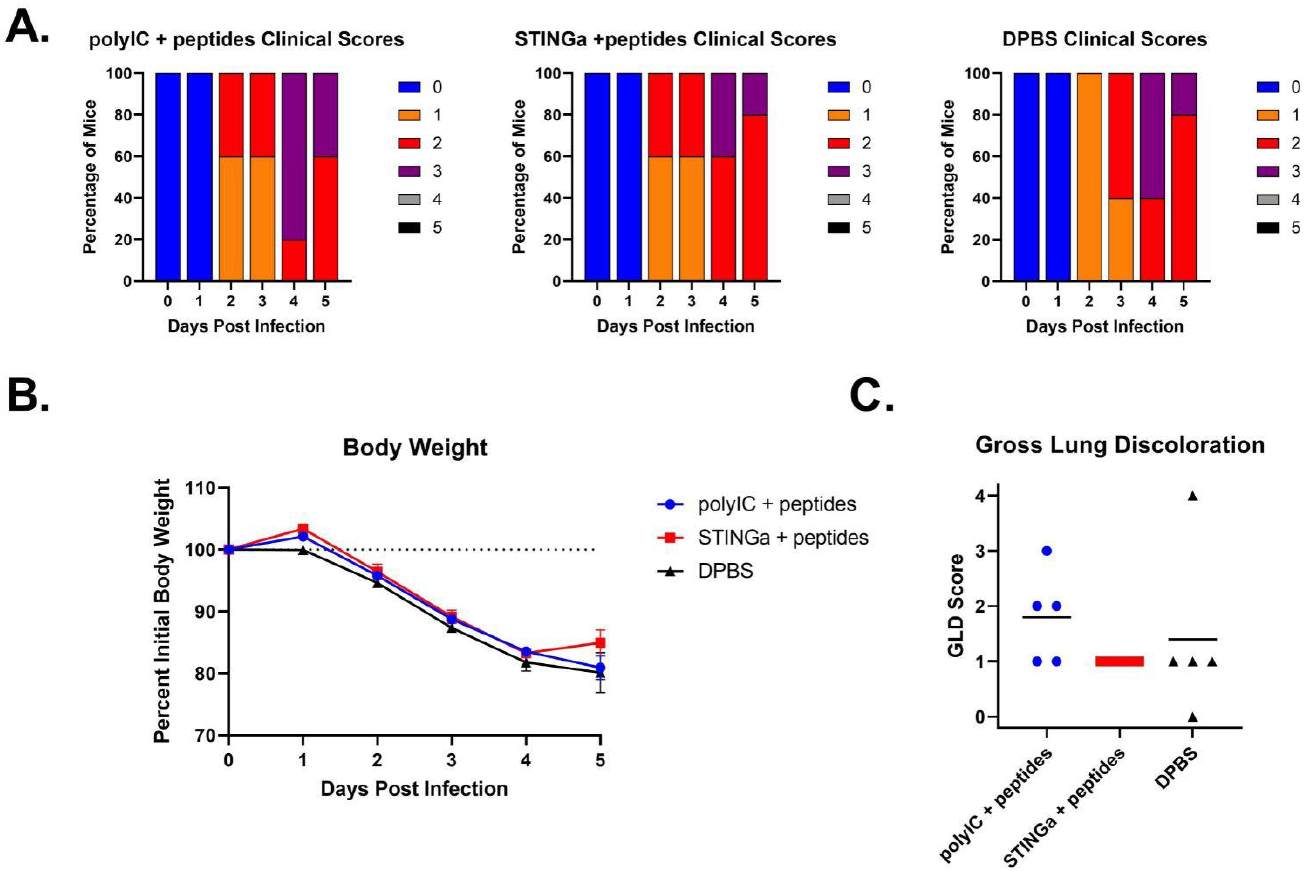
Lack of protection from infection or disease of BALB/C mice inoculated with SARS-CoV-2-MA10. Mice vaccinated with peptides and either poly(I:C) or STING agonist did not have statistically different clinical scores (A), loss of body weight (B) or gross lung discoloration (C) compared with mice given a control.

**Figure 5.**
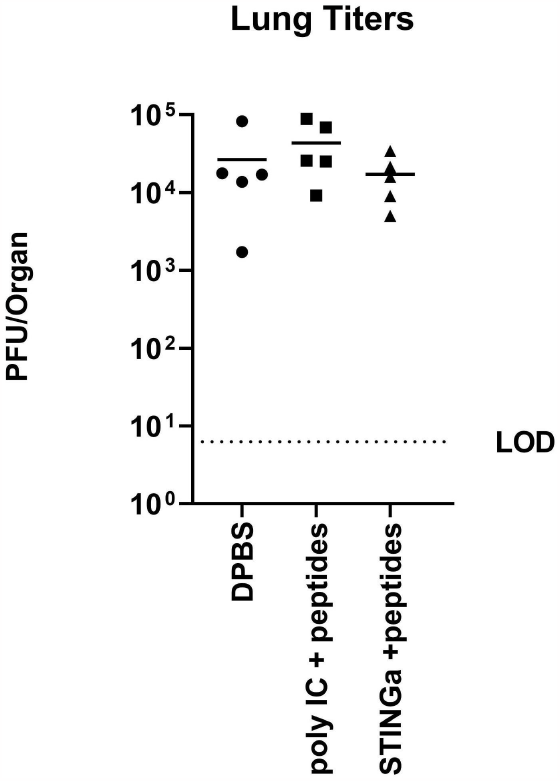
Viral titers in lungs of BALB/c mice after vaccination with one of poly(I:C) with vaccine peptides, STING agonist with vaccine peptides, or control, followed by challenge with SARS-CoV-2-MA10. There was no significant difference between groups.

## Discussion

We initially considered three distinct scenarios which could explain the lack of protection against severe disease despite strong virus specific T-cell responses in this study. The first possibility is that T-cell responses do not, in general, play any protective role against SARS-CoV-2 in the absence of antibody responses. In light of subsequent evidence which has emerged since this study was conducted, we now feel confident dismissing this explanation.

Another possibility is that in our particular model, BALB/c mice challenged with SARS-CoV-2-MA10, T-cell responses do not play any meaningful role in viral clearance independent of B-cells. Since related previous work has used different animals (e.g. K18-hACE2) and/or different viral strains (e.g. Wuhan-1), it is possible that lack of protection may be a peculiarity of our model. However, more recent studies continue to show evidence of T-cell protection against SARS-Cov-2 using diverse models. Thus, we now feel more confident in assigning a low probability to this explanation.

The last possibility is that there is significant mismatch between SARS-CoV-2 T-cell epitopes and those of BALB/c mice infected with SARS-CoV-2-MA10. Our vaccine peptides were primarily selected for predicted human immunogenicity. Though we did screen candidate peptides for predicted murine MHC binding, the murine filtering was less stringent than HLA binding predictions. Additionally, human T-cell epitope predictions were not limited to just HLA binding but also included a more comprehensive model of immunogenicity trained on human T-cell epitopes curated from IEDB(39). Though we cannot conclusively demonstrate this mismatch, we accept it as the most likely explanation for our negative findings.

